# Improving the sensitivity of fluorescence-based immunoassays by time-resolved and spatial-resolved measurements

**DOI:** 10.1101/2023.03.10.532014

**Authors:** Ran Kremer, Shira Roth, Avital Bross, Amos Danielli, Yair Noam

**Affiliations:** Faculty of Engineering, Bar-Ilan University, Max and Anna Webb Street, Ramat Gan, 5290002, Israel

**Author notes:** These authors have contributed equally to this work (Amos Danielli, Yair Noam).

**Keywords:** immunoassays, magnetic beads, image processing, signal processing, in vitro diagnostics

## Abstract

Detection of target molecules, such as proteins, antibodies, or specific DNA sequences, is critical in medical laboratory science. Commonly used assays rely on tagging the target molecules with fluorescent probes. These are then fed to high-sensitivity detection systems. Such systems typically consist of a photodetector or camera and use time-resolved measurements that require sophisticated and expensive optics. Magnetic modulation biosensing (MMB) is a novel, fast, and sensitive detection technology that has been used successfully to detect viruses such as Zika and SARS-CoV-2. While this powerful tool is known for its high analytical and clinical sensitivity, the current signal-processing method for detecting the target molecule and estimating its dose is based on time-resolved measurements only.

To improve the MMB-system performance, we propose here a novel signal processing algorithm that uses both temporally and spatially resolved measurements. We show that this combination significantly improves the sensitivity of the MMB-based assay. To evaluate the new method statistically, we performed multiple dose responses of Human Interleukin 9 (IL −8) on different days. Compared to standard time-resolved methods, the new algorithm provides a 2-3 fold improvement in detection limit and a 25% improvement in quantitative resolution.

## 1 Introduction

Target molecule detection, such as proteins, antibodies, or specific DNA sequences, within a population of other molecules, is critical in medical laboratory sciences. A typical molecule detection system consists of three elements. The first is a biological recognition component capturing the target molecule. The second is a reporting element, such as a fluorescent dye, quantum dots, or gold nanoparticles, which translates the biorecognition event into an analytically valuable signal. The third is a detector for the physical signal [1]. Due to their high sensitivity [2–4] and multiplexing [5], optical sensing techniques [2, 3, 5, 6] are the backbone of clinical diagnostic devices.

Optical detection systems usually consist of a photodetector [2, 7–9], such as a photomultiplier tube (PMT) or an avalanche photodiode (APD), or a camera [4, 10, 11], e.g., a charged coupled device (CCD) or a complementary metal-oxide semiconductor (CMOS). A camera image provides two-dimensional spatial information but is less effective when the physical phenomenon signal is weak and embedded in background noise. On the other hand, time-resolved measurements increase optical detection sensitivity, particularly in detecting and identifying single fluorescent molecules. These techniques usually use confocal fluorescent microscopy with a photodetector that detects the emission from a single fluorescent molecule when it transverses through a picoliter-size detection volume. This small detection volume significantly reduces the background noise originating from spurious fluorescence of impurities and Raman scattering of solvent molecules [12].

Time-resolved detection methods include fluorescence fluctuation spectroscopy (e.g., fluorescent correlation spectroscopy and fluorescence cross-correlation spectroscopy), single-molecule fluorescence burst detection [7], and pulse excitation with time-gated electronics [8]. While these methods are sensitive, they require the use of expensive and sophisticated optics and, thereby, are less applicable in point-of-care applications. Moreover, they do not allow the utilization of spatial information via further processing, as in the case of a camera.

Camera-based detection is used in several clinical and research devices, such as the MagPix™ (Luminex Corp., Austin, TX) [10, 13] and SIMOA™ (Quanterix Corp., Lexington) [4]. However, these devices receive 2D spatial data from stationary target molecules. Therefore, data analysis relies only on signal intensity rather than spatial characteristics, such as the spatial distribution of light, the number of bead aggregates, and their size.

Recently, we presented a novel technology, termed magnetic modulation biosensing (MMB), and demonstrated its analytical and clinical sensitivity in various serological [14, 15] and molecular assay [16], including Zika and SARS-CoV-2. In an MMB-based assay, magnetic beads and fluorescently labeled probes are attached to the target analyte to form a “sandwich” assay [17–19]. An alternating external magnetic field gradient condenses the magnetic beads (and subsequently, the target molecules with the fluorescently labeled probes) to the detection volume. It sets them in a periodic motion, thereby enabling the removal of the background signal from the oscillating target signal without complicated sample preparation.

Initially, the MMB system detected the oscillating fluorescence signal with a photomultiplier and demodulated it with a lock-in amplifier [14]. Unlike other optical detection systems that measure the signal from individual beads, the MMB collects the fluorescent signal from multiple beads, resulting in a relatively strong signal. Under such an intense signal, it is possible to use a simple camera to analyze the intensity of oscillating fluorescent molecules rather than the expensive PMT. Consequently, in more recent versions of the MMB, the PMT was replaced by an ordinary camera. The camera is not only inexpensive, but also offers another key advantage: 2D information. However, the MMB has so far ignored this information and utilized only time-resolved information.

Here, we designed a novel signal processing algorithm that for the first time uses both temporal and spatial resolved information. The algorithm significantly improves the sensitivity of the MMB system. To evaluate the new algorithm, we used the MMB-based interleukin-8 (IL-8) assay and conducted four experiments, each on a different day. The results show that the new algorithm provided 2–3-fold improvement in the limit of detection (LoD) while either maintaining the same or improving the quantitative resolution.

## 2 Materials and Methods

### 2.1 System Description

The camera-based MMB system (cf. [16, 20]) includes a 532 μm laser diode (CPS532, Thirlabs) working at 0.25 mW that generates a 3 mm diameter beam. The system also includes a dichroic beam splitter (Di02-R532, Semrock) that directs the beam onto an objectives lens focused to a 150 um beam size on a rectangular sample cell (W2540, Vitrocom) containing the fluorescently labeled probes, target molecules, and magnetic beads. Two electromagnets generate an alternating magnetic field gradient at 1 Hz, which pulls the magnetic beads and concentrates them into a small detection area, thus increasing fluorescence-detection sensitivity. The oscillating magnetic-field gradient drives the aggregated beads from side to side, in and out of a laser beam, thus enhancing the signal emitted from fluorescence molecules bound to the beads compared to the background noise induced by those unbound. The resulting signal amplitude, which is now more distinct, is proportional to the concentration of target molecules [17].

### 2.2 Interleukin-8 binding assay

We used conjugated magnetic beads (6.5 μm) (Bio-Plex Pro Human Chemokine IL-8 / CXCL8 Set #171BK31MR2, BioRad, Hercules, CA, USA) for the binding assay. We photobleached the beads for two hours, according to Roth et al. [21]. We used a 96-well plate, wherein each well mixed 50 μL of X2 photobleached conjugated magnetic beads with 50 μL solutions at different concentrations of a recombinant human Interleukin-8 (IL-8) protein (#574202, BioLegend, San Diego, CA, USA). The overall well concentration levels were 0.05, 0.1, 0.2, 1, 2, 5, 20, and 200 ng/L. After one hour of incubation, we added 50 *μ*L of detection antibodies (X1) to each well and further incubated it for 30 minutes. We then added 80 μL of X1 streptavidin phycoerythrin SA-PE (Bio-Plex Pro Reagent Kit III #171304090M, BioRad, Hercules, CA, USA) and incubated it for another 20 minutes. We made all incubations at room temperature on a rotator. Finally, we washed the magnetic beads once by placing the 96-well plate on a MagJET separation rack (MR02, Thermo Fisher Scientific, Massachusetts, USA) for four minutes, taking out the solution, and pipetting the beads with 200 μL of an assay buffer (PBS X1, 1% BSA w/v, 0.05% Tween-20). The washed beads were placed again on the separation rack, and the buffer was replaced with 100 μL of assay buffer for MMB measurements.

To have sufficient data for statistical analysis, we followed the protocol [22], where each trial reads the MMB output for a given IL8-protein concentration or blank sample [23]. The analysis includes four experiments comprising eight independent blank samples and four independent samples for each IL-8 concentration. The overall data includes 30 twelve-second video streams corresponding to blank samples and between 13-15 video streams for each other concentration. Figure 2 presents snapshots of an instance where the fluorescence molecule bulk is under the laser beam, dubbed on-frames, for various concentrations.

**Fig 1.**
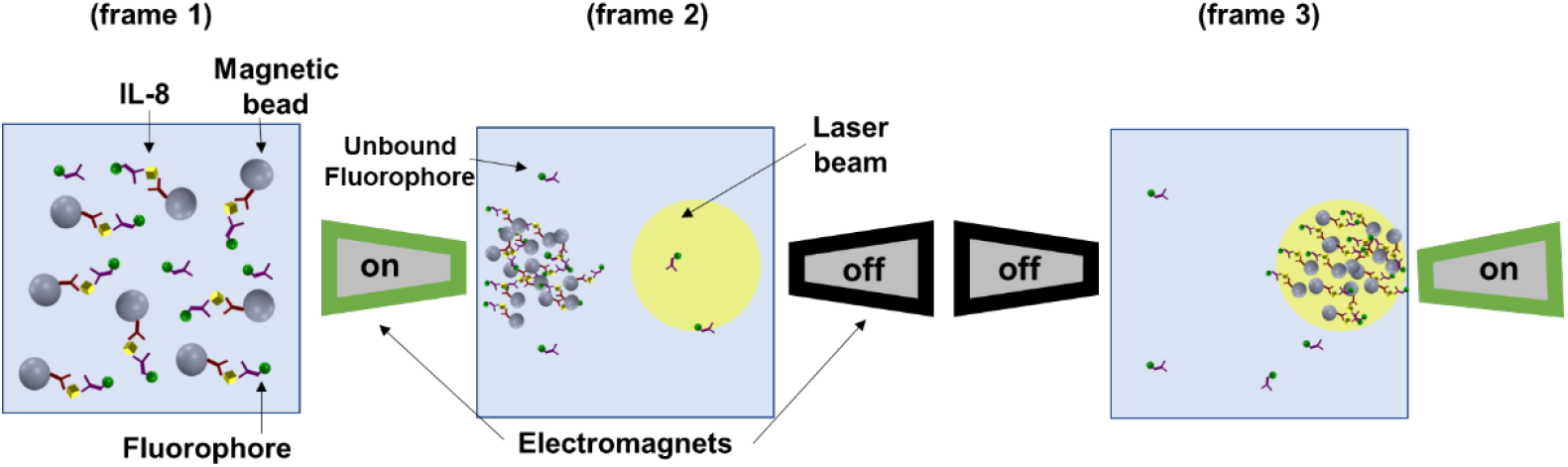
MMB operation modes: Frame 1 depicts Magnetic beads attached to IL-8 protein and detection antibodies while dispersed in the sample cell. Frame 2 depicts the state after applying an alternating magnetic field gradient that aggregates the magnetic beads from the entire sample to the detection volume. Unbound fluorophores stay dispersed in the sample cell. When the left electromagnet is “on,” the magnetic beads are aggregated to the left side of the sample cell, and the background signal emitted is recorded (frame 3). When the right-hand electromagnet is “on,” it modulates the magnetic beads to the right side of the sample cell inside the laser beam, and the emitted fluorescence is recorded.

**Fig 2.**
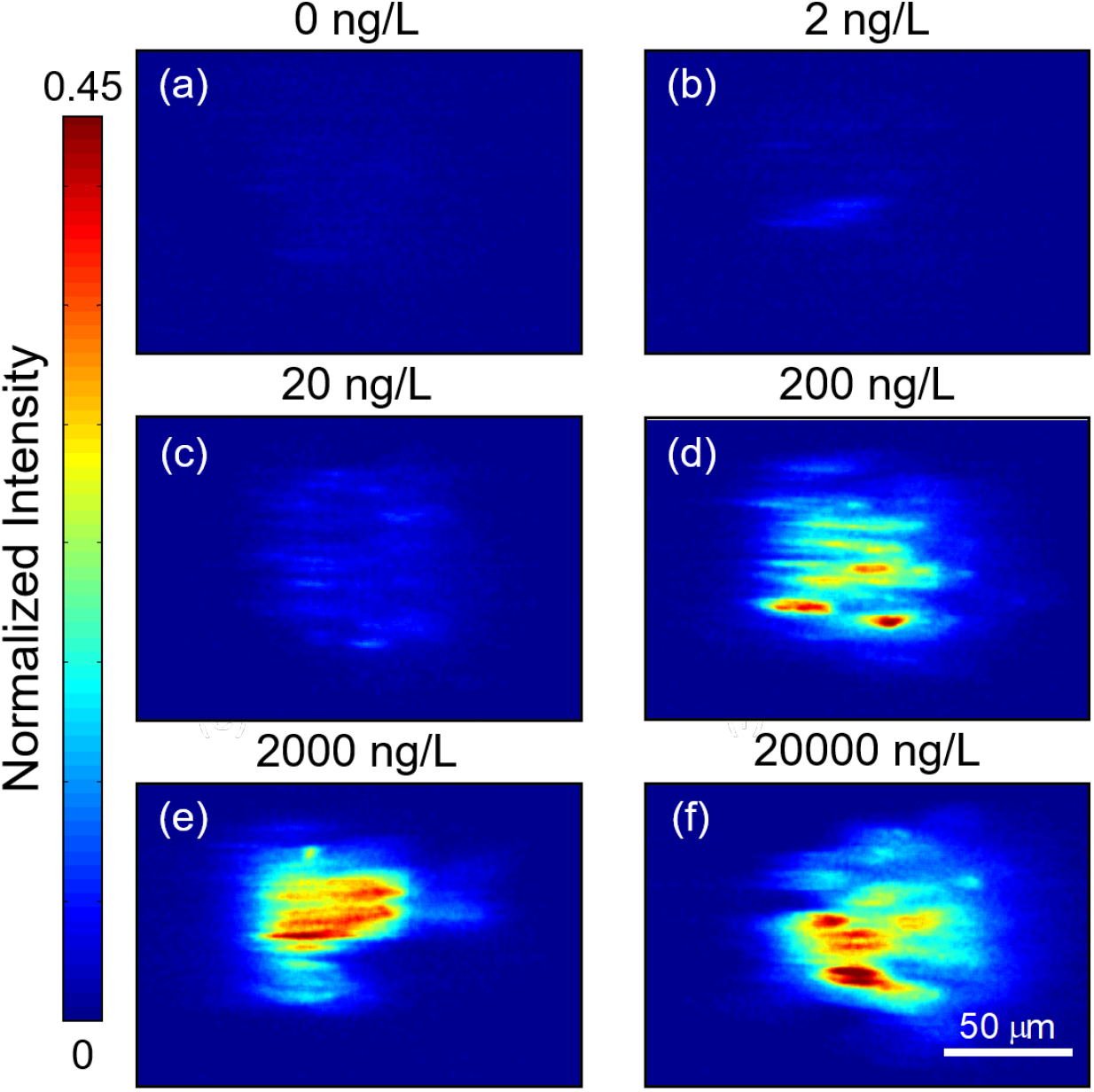
Snapshots at various IL-8 concentrations: (a) 0 ng/L, (b) 2 ng/L, (c) 20 ng/L, (d) 200 ng/L, (e) 20,000 ng/L, and (f) 200,000 ng/L.

#### 2.2.1 Mathematical Model

The MMB system generates a twelve-second video frame for each experiment at 50 frames-per-second (fps). We denote each video by 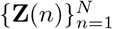, *N* = 50 × 12, where each 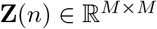 is a matrix representing a single video frame (image) at time instance *n* and **Z***_i,j_* (*n*) denotes its (*i, j*) entry. The latter corresponds to the light intensity impinging upon the corresponding camera pixel, which increases with the target-molecule concentration. Mathematically, consider a solution with a given concentration *c*, then each video frame satisfies

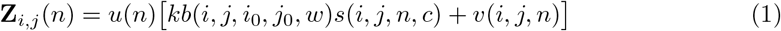

where *s*(*i, j, n, c*) indicates the fluorescence molecule concentration in the point corresponding to pixel-(*i, j*) at time instance n. Furthermore, b(·) denotes the laser spatial-signature (normalized), (*i*_0_, *j*_0_) marks the laser-beam center, and *k, w* are constants representing the laser intensity and decay rate, respectively. Finally, *u*(*n*) represents the bleaching effect, typically expressed by

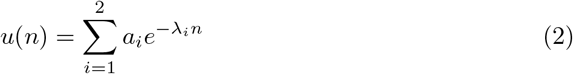

and *v*(·) denotes a spatial-temporal noise, which is mainly due to autofluorescent light emitted from the beads [18]. The magnetic modulation separates the signal from the background noise of the non-magnetized solution hence improving measurement sensitivity.

An aggregate is a bulk of fluorescently labeled target molecules, which allows sensitive biomarker detection even at low concentrations. We expect a noticeable aggregate in high target molecule concentrations, whereas, at intermediate and low concentrations, we expect dotted and blurred aggregates, respectively. The magnetic-field gradient is modulated at a frequency of 1 Hz [17].

The aggregate traverses the area illuminated by the laser beam at each cycle, thus inducing a different spatial signature. When the aggregate is entirely inside the laser beam, the image is brighter than when the aggregate is outside. The resulting signal is periodic at one Hz, with the intensity varying from high to low at each cycle. The camera captures 50 frames per second (fps), with multiple of these at each period. When the aggregate is inside the laser beam, we refer to the images as “on-frames,” whereas when it is outside, we use “off-frames.” Each video frame contains spatial information. For example, the on-frames have different spatial structures depending on the target molecule concentration. These include solid bulks at high concentrations while dotted and blurred at medium to low concentrations. Here, we use this information to improve the analytical performance of the MMB technology.

## 3 Signal Processing Algorithms for Enhanced MMB Performance

This section presents two novel signal-processing algorithms for enhancing MMB performance. Each algorithm corresponds to a feature with unique spatial and temporal processing. The first algorithm (Feature 1) improves both LoD and quantitative resolution (QR), whereas the second (Feature 2) improves only the LoD but more significantly than the former. The algorithm includes preliminary processing followed by a feature extraction phase. The preliminary processing reduces the frame size and detects the laser beam. We then use spatial processing to transform 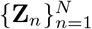 into a one dimensional time-series *y(*n*),* dubbed temporal-feature. Finally, we apply time-domain signal processing to extract the scalar feature *x* that constitutes the response.

### 3.1 Preliminary Processing

Here we reduce the frame size and compute the pinhole mask, which is necessary for the subsequent phase. We begin with detecting the laser-beam center, for which we average the video frames 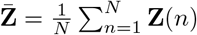. We then calculate 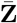 center of mass (*z_x_, z_y_*), where the horizontal coordinate is

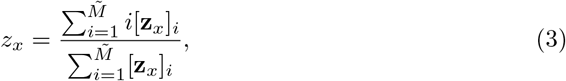

in which 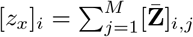. Similarly, we calculate the vertical coordinate *z_y_* by substituting **z***_y_* for **z***_x_* in (3), where 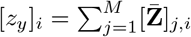. Then from each frame **Z**(*m*), we extract a 300 × 300 frame 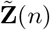 centered at (*z_x_, z_y_*); i.e.,

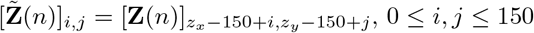. We detect the laser beam by reducing each frame to a tinier one, focusing on the region of interest centered at that beam. We apply a pinhole mask, a small circular aperture with a radius of 150, transferring only the central bright spot. Explicitly, let *R*_U_ be the pinhole radius, then the pinhole mask **P** is the following matrix

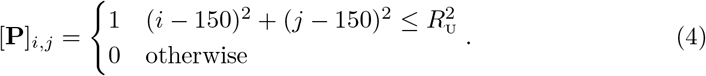

### 3.2 Feature Extraction

The data is high dimensional. Therefore, we represent it more compactly while preserving meaningful dose information; a process known as feature extraction. Here, we extract a scalar feature from every frame using spatial filtering. These features form a time series that is then processed to produce a dose estimator. In what follows, we describe these steps rigorously.

#### Feature 1 (Reference Interval Median (RI-M))

We begin with feature *x*_1_, dubbed RI-M. We first transform each frame **Z**(*n*) into 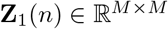 by extracting the pixels corresponding to the reference-interval within the 80th to 99.5th percentiles. Explicitly, let *α*_U_, *α*_L_ be the limits of the 99.5th and 80th percentiles, respectively, then

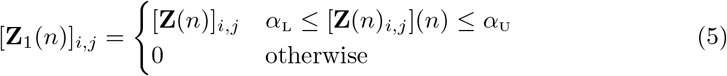

The resulting video 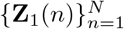 is transformed into a one-dimensional time-series 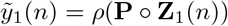, where **P** is the pinhole mask (4), ○ is the Hadamard product,^1^ and *ρ*(·) denotes the spectral matrix-norm; that is, *ρ*(**A**) is the largest singular value of the matrix **A**. We then turn to a one-dimensional signal processing of 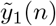 that applies a 12-length median filter

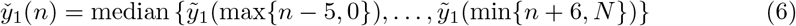

Finally we set

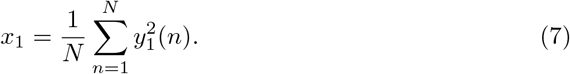

#### Feature 2 (Second SVD, Median Band Pass (SVD-M-BP))

Here *y*_2_(*n*) is the 2nd singular value in the singular value decomposition (SVD) of **Z**(*n*). Explicitly, let

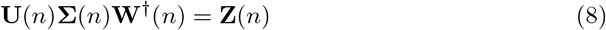

be the SVD of **Z**(*n*), where 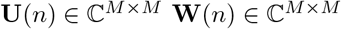 are unitary matrices and 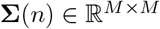 is a diagonal matrix containing the singular values *σ_m_*(*n*), *m* = 1,…, *M* in a decreasing order; that is, [**Σ**(*n*)]*_ij_* = *δ_i-j_σ_i_*(*n*) with *σ*_1_(*n*) ≥ *σ*_2_(*n*) ≥… *σ_M_*(*n*) ≥ 0, *δ_i_* is the Kronecker delta. Then, we set *y*_2_(*n*) = *σ*_2_(*n*).

Having obtained the one-dimensional signal, we now use a time-domain filter with three equal passbands of Δ*f* = 0.5 Hz centered at 1, 3, and 5 Hz, respectively. Let *h*_BF-3_(*n*) be the filter impulse response; the filter output is then

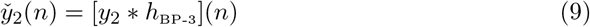

where [*x * y*](*n*) denotes the linear convolution between *x*(*n*) and *y*(*n*). The rationale behind the filter is that it corresponds to the first three harmonies of the Fourier series of a square wave at 1 Hz, which represents the magnetic modulation of the MMB. Finally, we set

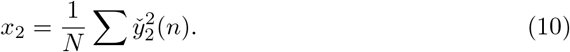

### 3.3 Baseline Approach

The baseline approach to which we compare the proposed algorithms, begins with preliminary processing (cf. Section 3.1). Then, we form a one-dimensional scalar time-series by mapping each video frame to its squared Frobenius norm, emulating the photomultiplier tube (PMT). We then apply a 12-length median filter followed by a 4 Hz cutoff low-pass filter. The final output is the resulting signal energy, denoted by *x*_BL_.

## 4 Results

This section compares the proposed algorithms to the baseline scheme for the experiment depicted in Section 2.1); a comparison that includes the dose-response, LoD, and QR for each method. We transformed each response using the natural logarithm to be homoscedastic and Gaussian. Moreover, we used Levene’s and Conover’s tests and Kolmogorov-Smirnov test to corroborate homoscedasticity and normality, respectively. The results are summarized in Tables 3 and 4 in Appendix A.

**Table 1.** LoB and LoD of the baseline approach and the proposed algorithms (Features 1 and 2). The table shows the LoD gain; that is, the new LoD divided by the baseline LoD. Also shown is the gain in percentage, that is, the baseline LoD minus the new one relative to the baseline LoD.

| Method | LoB [ng/L] | LoD [ng/L] | Gain | Gain (%) |
| --- | --- | --- | --- | --- |
| Baseline | 0.17 | 0.384 |  |  |
| Feature 1 | 0.082 | 0.212 | 1.8 | (44%) |
| Feature 2 | 0.042 | 0.158 | 2.4 | (58%) |

**Table 2.** Quantitative Resolution (QR) compared to the baseline approach. Also shown is the gain in percentage, that is, the QR gap relative to the baseline QR.

| Dose [ng/L] | Baseline QR [ng/L] | Feature 1 QR [ng/L] | Gain (%) | Feature 2 QR [ng/L] | Gain (%) |
| --- | --- | --- | --- | --- | --- |
| 1 | 0.27 | 0.19 | 29.6% | 0.24 | 11% |
| 2 | 0.61 | 0.53 | 13% | 0.64 | -5% |
| 5 | 1.57 | 1.17 | 25.4% | 1.44 | 8% |
| 20 | 6.73 | 4.99 | 25.8% | 6.9 | -2.5% |

**Table 3.** p-value for Kolmogorov-Smirnov test for Normality.

| Does<br>[ng/L] | Baseline | Feature 1 | Feature 2 |
| --- | --- | --- | --- |
| 0 | 0.73 | 0.30 | 0.17 |
| 0.05 | 0.83 | 0.22 | 0.29 |
| 0.1 | 0.09 | 0.49 | 0.18 |
| 0.2 | 0.88 | 0.34 | 0.86 |
| 1 | 0.51 | 0.40 | 0.60 |

**Table 4.** p-values for Levene’s and Conover’s tests.

| Test | Baseline | Feature 1 | Feature 2 |
| --- | --- | --- | --- |
| Levene | 0.08 | 0.46 | 0.67 |
| Conover | 0.12 | 0.56 | 0.94 |

For comparison, because every feature ranges over a different interval, we offset and normalize each feature into the interval [1, 2] via min-max normalization. Explicitly, let 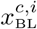 be the baseline response at concentration *c* ∈ *C* for sample *i* ∈ *I_c_*, where *I_c_* is the set of samples at *c*; then,

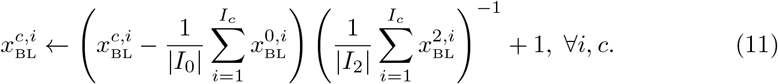

We repeated that procedure for the other features.

Figure 3 depicts the dose-response of the baseline approach. The limit of blank (LoB) and LoD are 0.17 and 0.384 ng/L, respectively. We used the guidelines in [22] with thresholds guaranteeing false positives and false negatives of no more than five %. Appendix A describes the method used to calculate each dose-response, and its LoB, LoD, and QR. Figure 4 depicts the dose-response of each feature and its corresponding LoB and LoD.

**Fig 3.**
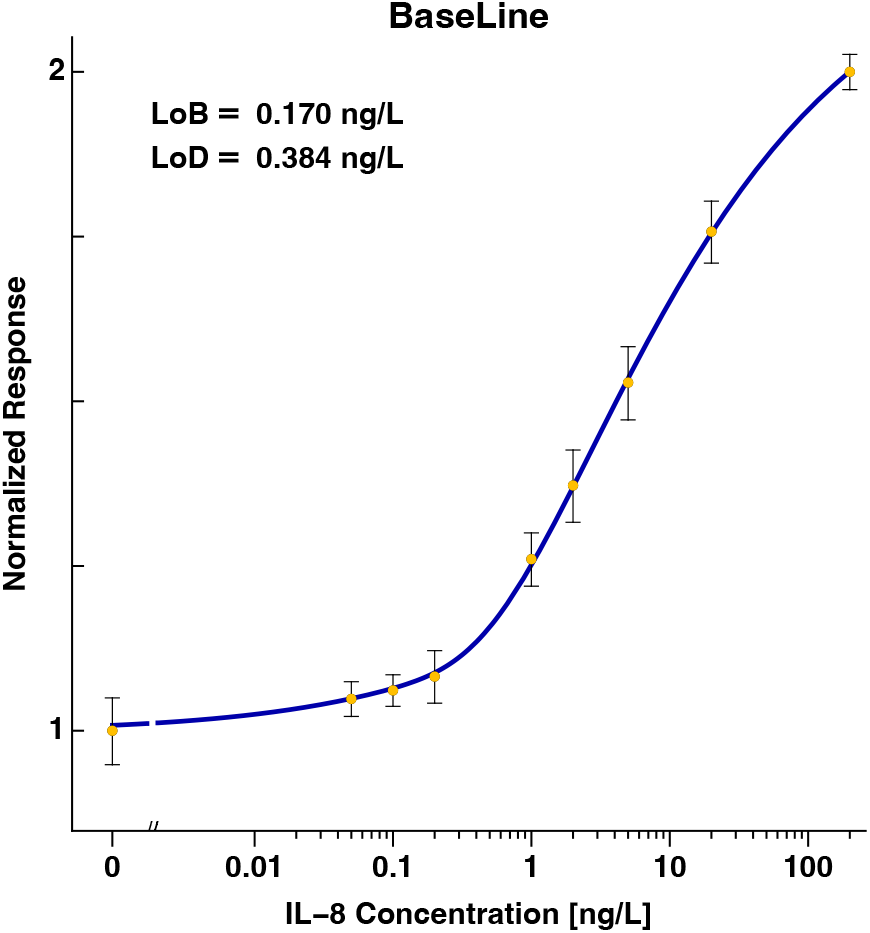
Dose response of the baseline approach.

**Fig 4.**
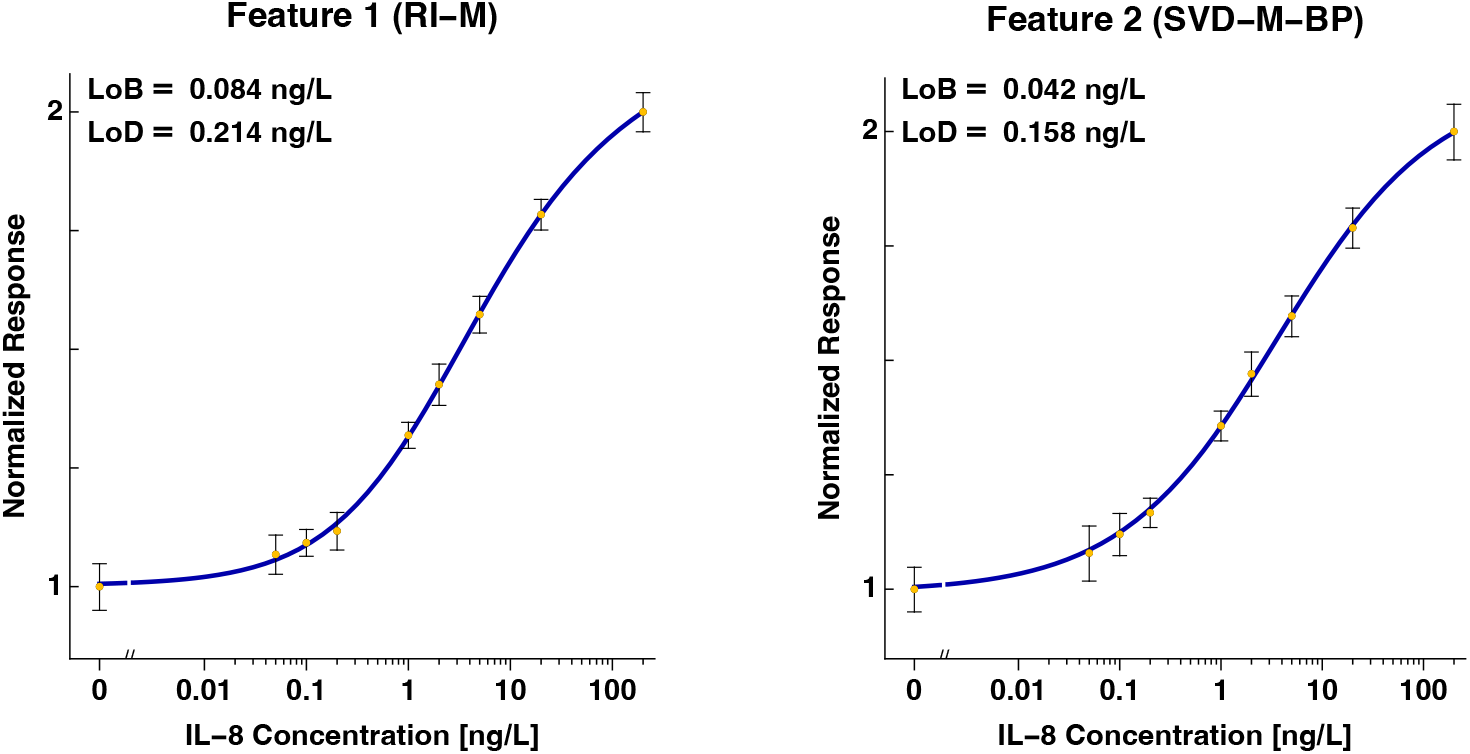
Dose responses Features 1 and 2 corresponding the proposed algorithms.

Table 1 summarizes the LoB and LoD of each method and the LoD gain compared to the baseline approach, whereas Table 2 summarizes the corresponding QR.

## 5 Discussion

To date, the MMB system has demonstrated fast detection and high analytical sensitivity in multiple serological and molecular assays, such as Zika [14] and SARS-CoV-2 [15, 16]. However, the signal processing included only time resolved measurements (i.e., the “baseline” approach). Here, we introduced two algorithms (“Feature 1” and “Feature 2”) that improved LoD and and estimation error (QR). Figure 4 indicates that spatiotemporal processing significantly improves target molecule detection (LoD) and estimation error (QR). Table 1 shows that Feature 1 improves LoD substantially by a factor of 1.8 (44%), whereas Feature 2 enhances LoD even further by a factor of 2.4 (58%). Table 2 details the QR at each concentration of all methods. The results show that Feature 1 has an improved QR, whereas Feature 2 does not. This outcome implies that the LoD gain of Feature 2 is at the expense of QR.

These results can be explained by the attributes associated with each feature. Feature 1 captures most of the energy of each video frame, including a DC signal component. It does so due to the reference interval and the spectral norm used in the spatial processing and because it applies only the median filter in the time domain.

Feature 2 includes more subtle information because its spatial processing involves only the second singular value. That value bears less energy than the spectral norm (the first singular value). Moreover, the time domain signal-processing of Feature 2 also includes the triple BPF, further reducing the noise. However, this noise reduction eliminates the DC signal. It also reduces other spectral components residing outside that filter narrow passbands at 1, 3, and 5 Hz. ^2^ The output signal, however, bears information beyond these bands since it is not a pure square wave embedded in noise. That output includes a DC component and other factors, such as an uneven duty cycle.

Due to the enhanced noise reduction, Feature 2 is more suitable for low-level detection than Feature 1. However, because of its improved QR, Feature 1 outperforms Feature 2 in higher concentrations. Hence, Feature 2 signal distortion outweighs its noise reduction in these concentrations.

## 6 Conclusions

This research proposed a new computational scheme for detecting and estimating target molecules within a molecular assay by the MMB system. For statistical assessment, we conducted four experiments; each repeated the same procedure at the same concentrations but on a different day. The algorithm has two operation modes; each corresponds to a different feature. The first feature improves detection performance (LoD) by a factor of 1.8 (44 %) while improving QR by 25% on average, in high concentrations. The second feature, aimed only at low lever detection, improves LoD by a factor of 2.4 (58%). These results demonstrate the effectiveness of spatiotemporal processing.

The results also pave the way for advanced machine learning methods to extract and combine additional features to enhance detection further. For example, we are looking at combining Features 1 and 2, each consisting of the first and second singular values. This fact suggests that the two features are uncorrelated; thus, performance can be improved if they are appropriately combined.

## Abbreviations

MMB: Magnetic modulation biosensing
QR: quantitative resolution
LoB: limit of blank
LoD: limit of detection
LPF: low pass filter
SVD: singular value decomposition

## Appendix

### A Dose-Response

#### A.1 Homoscedasticity and Normality

We used the natural logarithm function to transform the responses to be Normal and homoscedastic at low concentrations up to four times the LoB. Table 3 details the p-values of the Kolmogorov-Smirnov test for each method. The results corroborate that the data is Normal. We then tested the data for homoscedasticity using Levene’s and Conover’s tests. The results, detailed in Table 4, corroborate homoscedasticity. After corroborating Normality and homoscedasticity, we used the Cedergreen-Ritz-Streibig model [24] to fit the dose-response.

#### A.2 Levels of Blank and Detection

We determined the LoB and LoD via parametric tests [22]. For the LoB we calculated the critical signal ^3^ value (cf. [25], Sec. 3.7.4) corresponding to *α* = 5 % using

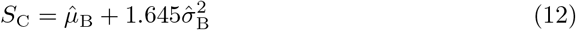

where 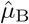 is the sample mean, and 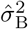 is the sample variance. We then determined the LOB (in the concentration domain) from *SC* via the inverse dose-response.

For the limit of detection, we estimated the pooled variance at concentrations up to four times the LoB, which we had already corroborated to be homoscedastic. Explicitly, for Features 1 and 2, we use 0.05, 0.1, and 0.2 ng/L, whereas the baseline further included 1 ng/L. Hence, the minimum detectable signal value is

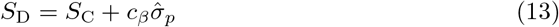

where *σ_p_* is the root pooled-variance, 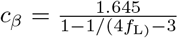 and *N*_L_ = 43 – 3 = 40. Finally, we set the LoD from *S*_D_ by via the inverse dose-response.

## Acknowledgments

This research was partially supported by the Israel Science Foundation (grant No. 1142/15).

## Author Contributions

**Conceptualization:** Yair Noam, Amos Danielli

**Data curation:** Ran Kramer, Yair Noam

**Formal analysis:** Ran Kramer, Yair Noam

**Funding acquisition:** Amos Danielli

**Investigation:** Shira Roth

**Validation:** Shira Roth, Ran Kramer

**Methodology:** Yair Noam, Amos Danielli

**Project administration:** Yair Noam

**Software:** Ran Kremer, Yair Noam

**Supervision:** Yair Noam, Amos Danielli

**Resources:** Yair Noam Amos Danielli

**Visualization:** Ran Kremer, Yair Noam, Shira Roth

**Writing – original draft:** Ran Kremer, Amos Danielli, Yair Noam

**Writing – review & editing:** Ran Kremer, Shira Roth, Yair Noam, Amos Danielli

## Footnotes

1 The Hadamard product is defined by the entry-wise product of two matrices.

2 These passbands correspond to the modulating signal of the MMB; that is, the 1 Hz square wave with a duty cycle of one-half, induced by the alternating magnetic field (cf. Section 2.1).

3 The term “signal” refers to the feature used to derive the dose-response.

## Notes

### Competing Interest Statement

The authors have declared no competing interest.

